# Functional diversity of PFKFB3 splice variants in glioblastomas

**DOI:** 10.1101/2020.10.09.332817

**Authors:** Ulli Heydasch, Renate Kessler, Jan-Peter Warnke, Klaus Eschrich, Nicole Scholz, Marina Bigl

## Abstract

Tumor cells tend to metabolize glucose through aerobic glycolysis instead of oxidative phosphorylation in mitochondria. One of the rate limiting enzymes of glycolysis is 6-phosphofructo-1-kinase, which is allosterically activated by fructose 2,6-bisphosphate which in turn is produced by 6-phosphofructo-2-kinase/fructose-2,6-bisphosphatase (PFK-2/FBPase-2 or PFKFB). Mounting evidence suggests that cancerous tissues overexpress the PFKFB isoenzyme, PFKFB3, being causing enhanced proliferation of cancer cells.

Initially, six PFKFB3 splice variants with different C-termini have been documented in humans. More recently, additional splice variants with varying N-termini were discovered the functions of which are to be uncovered.

Glioblastoma is one of the deadliest forms of brain tumors. Up to now, the role of PFKFB3 splice variants in the progression and prognosis of glioblastomas is only partially understood. In this study, we first re-categorized the PFKFB3 splice variant repertoire to simplify the denomination. We investigated the impact of increased and decreased levels of PFKFB3-4 (former UBI2K4) and PFKFB3-5 (former variant 5) on the viability and proliferation rate of glioblastoma U87 and HEK-293 cells. The simultaneous knock-down of PFKFB3-4 and PFKFB3-5 led to a decrease in viability and proliferation of U87 and HEK-293 cells as well as a reduction in HEK-293 cell colony formation. Overexpression of PFKFB3-4 but not PFKFB3-5 resulted in increased cell viability and proliferation. This finding contrasts with the common notion that overexpression of PFKFB3 enhances tumor growth, but instead suggests splice variant-specific effects of PFKFB3, apparently with opposing effects on cell behaviour. Strikingly, in line with this result, we found that in human IDH-wildtype glioblastomas, the PFKFB3-4 to PFKFB3-5 ratio was significantly shifted towards PFKFB3-4 when compared to control brain samples. Our findings indicate that the expression level of distinct PFKFB3 splice variants impinges on tumorigenic properties of glioblastomas and that splice pattern may be of important diagnostic value for glioblastoma.

## Introduction

Glioblastoma is the most common malignant primary tumor in brain. The high rate of aerobic glycolytic flux, a mechanism known as the Warburg effect, is a metabolic hallmark of tumors including glioblastoma [1]. As a result, glioblastoma cells possess increased levels of fructose-2,6-bisphosphate (F2,6BP), the main regulator of 6-phosphofructo-1-kinase, which in turn represents one of the rate-controlling glycolytic enzymes [2, 3]. Both synthesis and degradation of F2,6BP are catalysed by 6-phosphofructo-2-kinase/fructose-2,6-bisphosphatase (PFK-2/FBPase-2, in human PFKFB, EC 2.7.1.105/EC 3.1.3.46), which belongs to a family of homodimeric bifunctional enzymes [4]. In human, there are four major PFKFB isoenzymes encoded by four genes (*PFKFB1-4*), which possess high sequence homologies within their catalytic core domains. PFKFB isoenzymes differ in pattern and level of expression as well as in functional properties including their response to protein kinases [5]. Typically, PFKFBs have a similar capacity to function as kinase and bisphosphatase. However, for PFKFB3 this balance has been shown to be shifted towards kinase activity, which in turn enables sustained high glycolysis rates [6]. *PFKFB3* gene is localized on chromosome 10p15.1 [7] and is ubiquitously distributed throughout human tissues. It shows elevated levels in rapidly proliferating cells such as tumorigenic and leukemic cells [8]. Both inflammatory and hypoxic stimuli were shown to trigger PFKFB3 expression [9, 10]. Consistently, *PFKFB3* contains multiple copies of the oncogene-like AUUUA instability element within its 3’ untranslated region [7]. Moreover, PFKFB3 was found to be shuttled to the nucleus by a process which appears to be triggered by a highly conserved nuclear localization motif within the C-terminus [11]. F2,6BP synthesized in the cell nucleus increases cyclin-dependent kinase (CDK)–dependent phosphorylation of the CIP/KIP-protein p27, which is subsequently degraded in the proteasome [12]. PFKFB3 was also reported to participate in G2/M transition [13] and to regulate the cell cycle (transition from G1 to S phase) by binding to cyclin dependent kinase 4 (CDK4) [14]. Gustafsson *et al*. (2018) identified PFKFB3 as a critical factor in homologous recombination repair of DNA double-strand breaks [15]. Conclusively, PFKFB3 constitutes a metabolic key player, which causally couples cell cycle and glucose metabolism to proliferation of cancer cells [16]. In humans, six PFKFB3 splice variants (designated UBI2K1-6) have been described [17]. The diversity of these transcripts results from a combination of different exons encoding varying PFKFB3 C-termini (Fig 1). Splice variant UBI2K5 and its role in cancer metabolism was studied in detail [18, 19], but thus far the role of most other splice variants remains enigmatic. Kessler *et al*. [20] found increased expression levels of total PFKFB3 in high-grade astrocytomas compared to low-grade astrocytomas and non-neoplastic brain tissue. Healthy brains express the entire set of PFKFB3 splice variants (UBI2K1-6). In contrast, glioblastoma predominantly express UBI2K4-6 with UBI2K5 and UBI2K4 being increased and decreased respectively compared to tissue from control brains [21]. Based on this inverse correlation between UBI2K4 expression and the growth rate of cells, Zscharnack *et al*. [21] concluded that UBI2K4 suppresses tumor cell growth. To elucidate the impact of UBI2K4 on the metabolism of cancer cells in detail we analyzed UBI2K4 deficient HEK-293 and a glioblastoma cell line (U87) with respect to their viability and proliferation capabilities.

**Fig 1.**
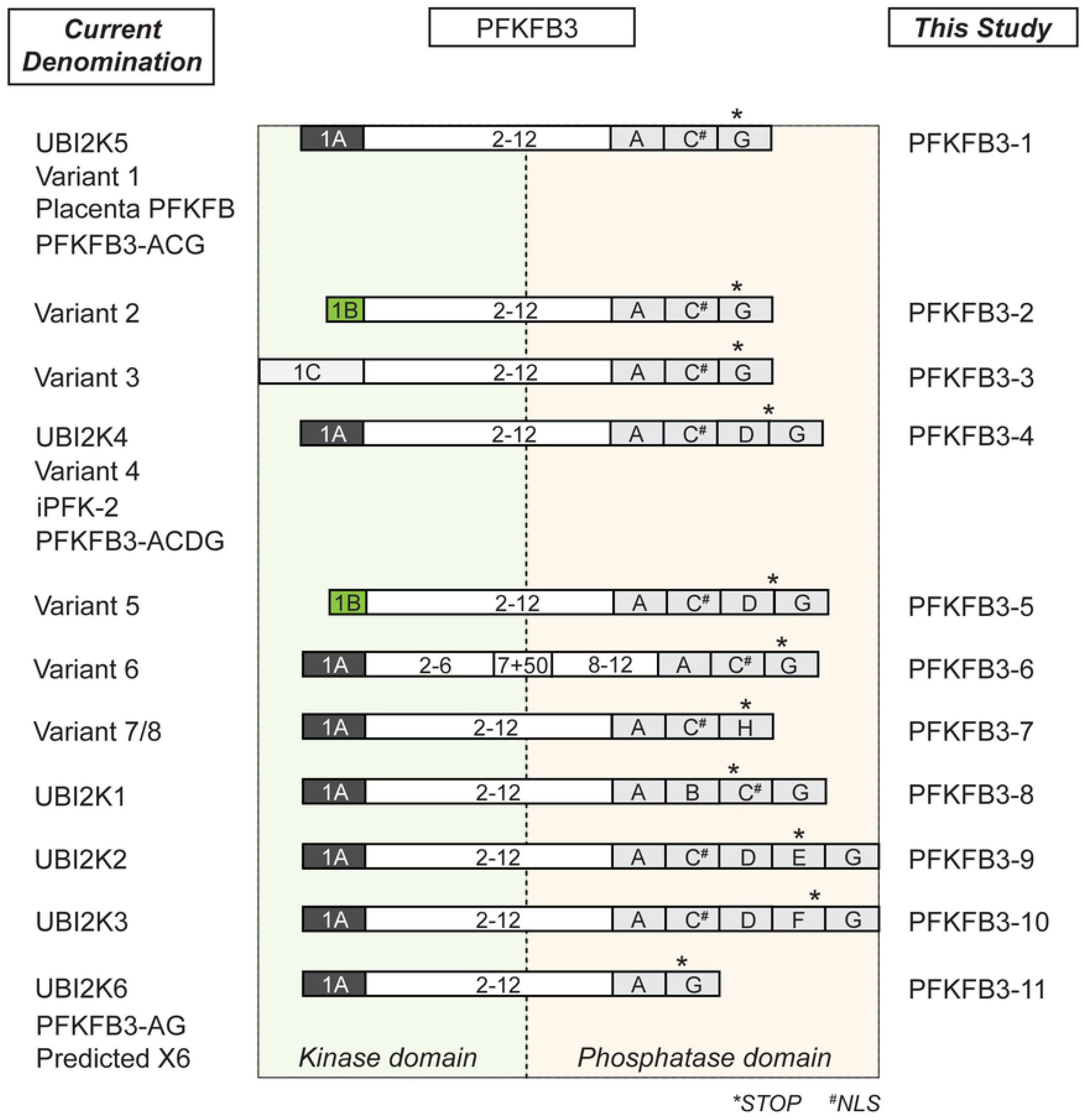
Schematic illustration of the transcript repertoire generated from the human *PFKFB3* gene locus. Schemes based on sequence analyses carried out using Basic Local Alignment Search Tool (BLAST) and Multalin interface page (multalin.toulouse.inra.fr/multalin). Left panel shows the current denomination of splice variants in literature and NCBI database; right panel indicates the denomination PFKFB3-1-11 used in this study. Conserved exons are depicted as white boxes with numbers except for PFKFB3-6, which contains an additional insert of 50 bp in exon 7. Variable N-terminal and C-terminal exons are colored and indicated by capital letters. * indicates the stop codon of each splice variant, # indicates the predicted nuclear localization signals (NLS).

In the past, the denomination of different PFKFB3 splice variants differed across laboratories. As a result, identical isoforms are often non-uniformly referenced. For example, the predominant splice variant in human brain (UBI2K5) is referred to as the ubiquitous PFK-2/FBPase-2 [22], placenta PFK-2/FBPase-2 [23] “Progestin Responsive Gene 1” [24] and PFKFB3-ACG [18] despite identical amino acid sequences of the respective proteins. Similarly, UBI2K4 and inducible PFK-2 (iPFK-2) [25] refer to the same molecule. To confuse matters even more, recently NCBI-PubMed published additional PFKFB3 splice variants designated ‘variant 1–7 and also putative splice variants designated as X2-X8’. Two of them are synonyms for previously described splice variants UBI2K4 (variant 4) and UBI2K5 (variant 1) and also the putative splice variant X6 is synonymous for UBI2K6. Variant 5 closely resembles UBI2K4 (variant 4), however, both differ in their N-termini. The present paper focuses on investigating in detail the impact of UBI2K4 (variant 4) and variant 5 on the metabolism of glioblastoma cells. Thus, to unambiguously refer to particular splice variants in this study we utilize a straight-forward PFKFB3 nomenclature with numbers referencing isoenzyme and splice variants (Fig 1). Therefore, variant 4 (UBI2K4) and variant 5 are designated as PFKFB3-4 and PFKFB3-5.

In this paper, we report a decreased viability and proliferation rate of PFKFB3-4 and PFKFB3-5-deficient U87 and HEK-293 cells, which was accompanied by a reduction in colony formation. Overexpression of PFKFB3-4 but not PFKFB3-5 resulted in increased cell viability and proliferation. In IDH-wildtype glioblastomas, the ratio of PFKFB3-4 to PFKFB3-5 was significantly shifted towards PFKFB3-4 compared to control brain samples. Our findings indicate different roles for splice variants PFKFB3-4 and PFKFB3-5 in healthy as well as malignant cells and implicate an important diagnostic role of these specific PFKFB3 splice variants in glioblastomas.

## Materials and methods

### Sample collection and genotyping

The study included 30 isocitrate dehydrogenase (IDH) -wildtype glioblastomas of World Health Organization grade IV, which were diagnosed as primary glioblastomas without clinical history. The glioblastomas were resected from patients undergoing neurosurgery at the Department of Neurosurgery, Paracelsus Hospital Zwickau (Germany). Histopathological and molecular diagnosis were done by K. Petrow (Institute of Pathology, Zwickau, Germany) and C. Mawrin (Department of Neuropathology, Otto-von-Guericke University, Magdeburg) based on the World Health Organization Classification [26]. The 15 surgical specimens of tumor-adjacent, macroscopically normal brain tissues according to criteria thoroughly described, were used as controls (Table S1). The ethics committee of the University of Leipzig approved this study (Reg. No. 167-14-02062014).

Genomic DNA of tumor samples was screened for IDH1 and IDH2 mutations as previously described Hartmann [27]. To analyze the IDH1 locus we used primers IDH1f and IDH1r, for IDH2 we used IDH2f and IDH2r (Table S2). Primers used in this study were synthesized by Metabion (Martinsried, Germany).

Unless stated otherwise, PCR products and plasmids generated in this study were sequenced using the BigDye Terminator Cycle Sequencing Kit and the Applied Biosystems 3130xl Genetic Analyzer (Applied Biosystems, Weiterstadt, Germany).

### Cell culture

The following cell lines were used: HEK-293 (ATCC CRL-1573); U87-glioblastoma cell line (ATCC HTB-14); SH-SY5Y (ATCC CRL-2266); 1321N1 human astrocytoma cell line (ECACC 86030402); LN-405 glioblastoma cell line (ACC 189). All cell cultures were maintained at 37°C in humidified atmosphere containing 5% CO_2_ and grown as monolayers in DMEM (Biochrom, Berlin, Germany), supplemented with 4.5 g/l glucose, 10% fetal bovine serum (Hyclone, Bonn, Germany), 1% penicillin/streptomycin/neomycin (Invitrogen, Karlsruhe, Germany). U87 and SH-SY5Y cells were grown in DMEM additionally supplemented with 1% non-essential amino acids (Invitrogen, Karlsruhe, Germany).

### RNA and protein isolation

Total RNA and protein from tissue and cell culture samples were extracted using TRIzol according to the manufacturer’s protocols (Invitrogen, Karlsruhe, Germany). Concentration and quality of RNA were determined by spectrophotometry using the NanoDrop^®^ ND-1000 (PeqLab, Erlangen, Germany). Total protein content was measured using the BioRad DC protein assay kit (Munich, Germany).

### Construction of shRNA-encoding plasmids for PFKFB3-4+5 silencing

Human PFKFB3-4+5 specific shRNA was designed as a 63-mer containing a hairpin-loop, which was cloned into H1 RNA polymerase promoter-containing pSuper vector. The vector contains an inducible system to stably integrate siRNA and an EGFP cassette [28]. A Zeocin resistance cassette was used to select stably transfected cells. For siRNA experiments, an overlapping sequence-fragment between exon D and G (Fig 2A) in the C-terminus of PFKFB-4 and PFKFB-5 was used. The shPFKFB3-4+5-coding sequences and sh-scrambled sequences are listed in Table S2. As judged from BLAST search scr-shRNAs show no significant sequence similarity to mouse, rat, or human gene sequences. The oligonucleotides were annealed and subcloned downstream of the H1 promoter into pTER-EGFP using *Hind*III and *Bgl*II.

**Fig 2.**
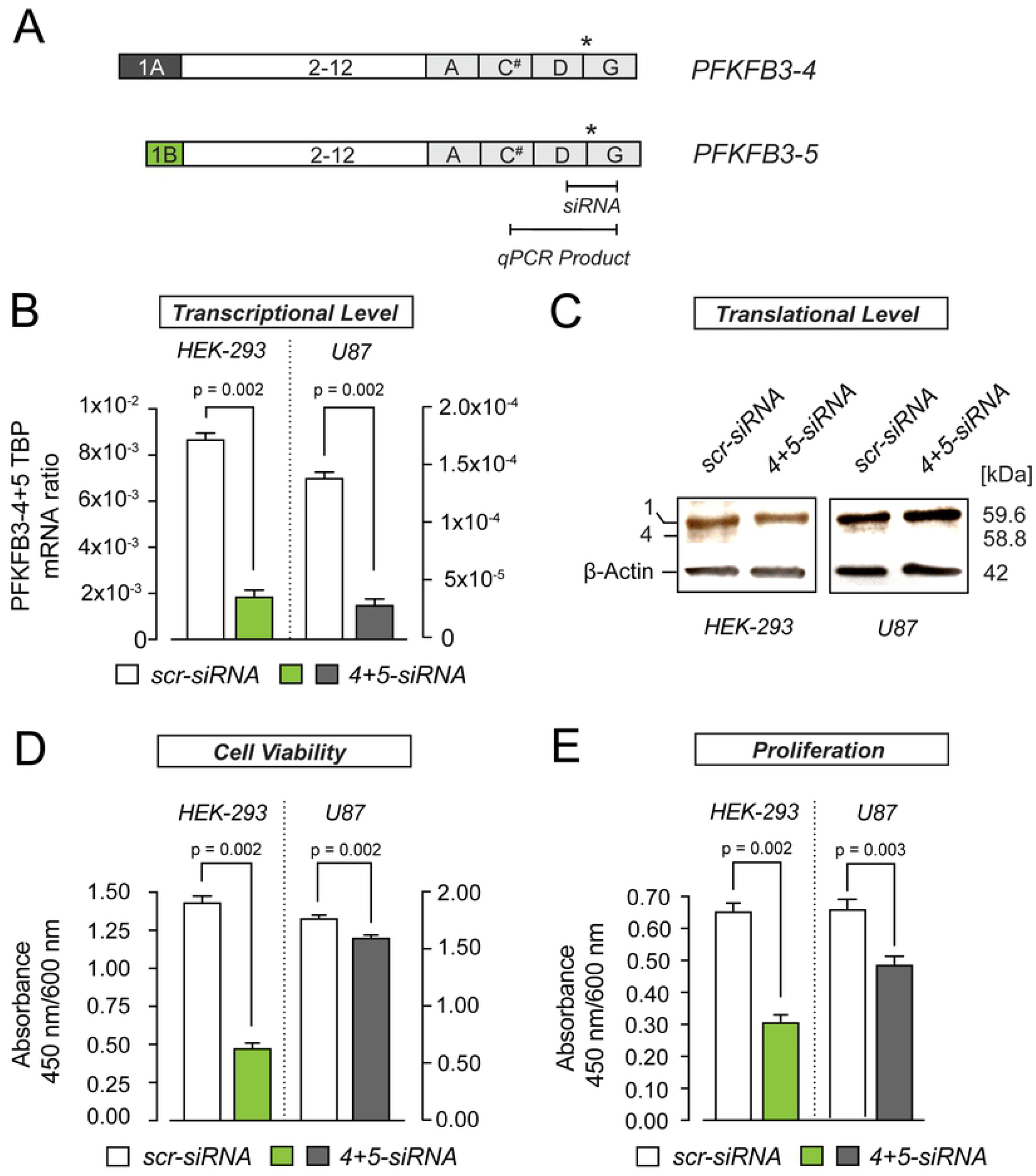
Knock-down of PFKFB3-4+5 alters proliferation and viability of HEK-293 and U87 cells. (A) Schematic representation of PFKFB3-4 and −5 with indication of PCR/qPCR as well as siRNA target sequences. Note that PFKFB3-4 and −5 differ in their N-but not C-termini. * indicates the stop codon, # indicates the predicted nuclear localization signal (NLS). siRNA probes used for gene silencing recognize sequences between exons D and G, in PFKFB3-4 and PFKFB3-5. (B) Quantification of stably and transiently inhibited PFKFB3-4+5 expression in HEK-293 and U87 cells, respectively. mRNA levels in cells carrying PFKFB3-4+5 siRNA was compared to scr-siRNA carrying cells. PFKFB3-4+5 expression was normalized to the amount of TBP mRNA measured by quantitative PCR. (C) Western blot analysis of PFKFB3 protein expression following siRNA mediated PFKFB3-4+5 knock-down utilizing polyclonal PFKFB3 antibody. β-Actin served as loading control, 30 μg protein per lane were applied. Western blot shows the 58.8 kDa band in scr-siRNA cells. The PFKFB3-5 protein was not detectable. (D) Quantification of cell viability of PFKFB3-4+5-depleted HEK-293 and U87 cells compared to scr-siRNA treated cells via WST-1 assay. (E) Quantification of cell proliferation of PFKFB3-4+5-depleted HEK-293 and U87 cells compared to scr-siRNA treated cells via BrdU-test. All values present the mean ± SEM from five independent experiments measured in duplicates (N=5, n=2).

### Engineering of PFKFB3-4 and PFKFB3-5 overexpression plasmids

Total RNA was obtained from astrocytoma cell line 1321N1 using TRIzol and reverse-transcribed with Transcriptor Reverse Transcriptase according to the manufacturer’s instructions (Roche Diagnostics, Mannheim, Germany). Full-length human PFKFB3-4 was generated by standard PCR using primers PFKFB3-4 reverse and PFKFB3-4 forward (Table S2). The resulting amplicon was cloned into pcDNA3.1/Hygromycin plasmid vector (Invitrogen, Waltham, MA, USA) using *Apa*I and *Afl*II. Similarly, the full-length fragment of human PFKFB3-5 was amplified with primers PFKFB3-5 reverse and PFKFB3-5 forward and subcloned into the plasmid pGEM-T using the T/A Cloning Kit (Promega, Mannheim, Germany). Subsequently, the amplicon was digested with *Xba*I and *Hind*III and inserted into the pcDNA3.1/Hygromycin plasmid vector (Invitrogen, Waltham, MA, USA). After confirmation by sequencing and enzymatic digest, both constructs were assigned the names pcDNA-PFKFB3-4 and pcDNA-PFKFB3-5.

### Generation of stable cell lines

Transfection of plasmids for overexpression purposes was performed using X-tremeGENE™ HP (Roche Diagnostics) according to the manufacturer’s instructions. To generate stably-expressing HEK-293 cell lines 150 μg/ml hygromycin B (Invitrogen) was added to the medium. Individual hygromycin-resistant colonies were selected and expanded.

Transfection of plasmids for knockdown purposes (shRNA-vectors) was performed using FuGene^®^HD (Roche Diagnostics) according to the manufacturer’s instructions. To generate stable cell lines the transfected cells were selected with Zeocin (200 μg/ml, Invitrogen, Waltham, MA, USA) and by EGFP fluorescence.

### Transient overexpression of PFKFB3-5

Transient transfection of plasmids for overexpression purposes was performed using X-tremeGENE™ HP (Roche Diagnostics) according to manufacturer’s instructions. The cells were harvested 24-48 hours after transfection. mRNA was measured 24 h after transfection and protein was measured 48 h after transfection.

### Transient knockdown in U87 cells

For transient knockdown of PFKFB-4 and PFKFB-5 in U87 cells, duplex siRNA was obtained from Thermo Fisher Scientific Biosciences (St. Leon-Rot, Germany) with UU overhangs (standard) The sequences are listed in Table S3.

Transfection for transient knockdown purposes (siRNA) was performed using DharmaFECT™ transfection reagents (Thermo Fisher Scientific) according to the manufacturer’s protocol. The transfection reagents were used with a final siRNA concentration of 25 nM. The cells were harvested 24-48 h after transfection (mRNA: 24 h, protein: 48 h).

### qPCR

For quantitative PCR, 500 ng of total RNA were reverse-transcribed using Transcriptor Reverse Transcriptase (Roche Diagnostics) and oligo-d(T)_n=18_ primer (Metabion) according to the manufacturer’s protocol. For quantification of PFKFB3-4+5, the cDNA was amplified in a LightCycler (Roche Diagnostics) using primers 3PFK2fo2 and iPFK2re6 as well as the LightCycler FastStart DNA Master Plus Set SYBR Green I Kit (Roche Diagnostics) according to the instruction manual.

To reliably calculate the RNA concentration, we generated RNA standards. To this end, a specific PFKFB3-4+5 fragment (Fig 2A) was reverse transcribed from total RNA of human brain (see Table S1, Pat.-No. 104) and PCR-amplified using primers 3PFK2fo2 and iPFK2re6 (Table S2). The PCR product was cloned into the pGEM-T vector. Sense strand RNA was transcribed using the Megascript *in vitro* Transcription Kit (Ambion, Wiesbaden, Germany) according to the manufacturer’s instructions to yield standard RNA. Standard curves were generated during each RT-PCR by serial fivefold dilution as previously described [21].

The TATA box binding protein (TBP) standard synthesis and the TBP quantification were carried out with primers TBPfo and TBPre. PFKFB-1 and PFKFB-11 were quantified as above with the primer pairs iPFK2Fo/PFK2Re and HBF10/6PFK2re5, respectively. RNA standards for PFKFB-1 and PFKFB-11 were synthesized as described for PFKFB3-4+5.

### Multiplex PCR

To pinpoint differences in the expression of PFKFB3-4 and PFKFB3-5, a multiplex PCR was established using PFKFB3-4 and PFKFB3-5 specific forward primers 4_Fo and 5_Fo as well as reverse primer 4/5_Re, which anneals to both splice variants (Fig 7A). 500 ng total RNA were reverse-transcribed with Transcriptor Reverse Transcriptase (Roche Diagnostics) using the primer 4/5_Re. PCR was performed using a master mix including the Expand high fidelity Taq polymerase (Roche Diagnostics). Amplicons were analyzed by standard agarose gel-electrophoresis. The ratio of PCR fragments was calculated from the intensity values of DNA bands analyzed with Herolab E.A.S.Y Plus Video gel documentation system (Herolab, Wiesloch, Germany).

To estimate the sensitivity of the primer pairs in the multiplex system, standard curves were established and the efficiency of the PCR was tested. The standard RNAs were synthesized from both target cDNA, which were subcloned in pGEM-T by an *in vitro* RNA synthesis kit (MAXIscript; Ambion). The copy numbers of RNA molecules were calculated on the basis of their absorbance values. The RNA products were serially diluted to prepare standard RNA solutions and were subjected to RT-PCR as described above.

### Western blotting

5-30 μg protein per lane were separated by standard SDS-PAGE (7,5% acrylamide gel) and semi-dry blotted onto nitrocellulose membranes (PALL Life Sciences, Dreieich, Germany).

The membranes were blocked with 5 % skimmed milk in Tris-buffered saline Tween 20 (TBST) for 2 h. For knockdown experiments the membranes were incubated with primary antibodies: rabbit-anti-human PFKFB3 (1:1000; ABIN 392768, Abgent/Biomol, Hamburg) and goat-anti-*β*-Actin IgG (1:5000; Santa Cruz Biotechnology, Heidelberg). For overexpression experiments PFKFB3 antibody (1:1000) and mouse-anti-*β*-Tubulin Antibody (1:5000; E7, DSHB, Iowa, USA) were used. Secondary antibody was incubated for 1 h at 25 °C with donkey-anti-rabbit IgG POD (1:30000, Dianova, Hamburg), donkey-anti-goat IgG POD (1:120000; Santa Cruz Biotechnology, Heidelberg) or goat-anti-mouse IRDye 800CW (1:15000; Li-COR, Nebraska, USA). Proteins were visualized using an enhanced chemiluminescence kit (SuperSignal West Dura, Thermo Fisher Scientific). To detect *β*-Tubulin the Odyssey FC 2800 (Li-COR Biosciences, Bad Homburg, Germany) was used.

### Cell viability and cell proliferation

Cell viability was evaluated using the colorimetric WST-1 assay (Roche Diagnostics). After a 4-h incubation period with WST-1 reagent the absorbance was measured at 450 nm/ 600 nm using a microplate reader (ELISA-Reader Zenyth 200st, Anthos, Krefeld, Germany).

Cell proliferation was evaluated using a colorimetric bromodeoxyuridine (BrdU) cell proliferation ELISA kit (Roche Diagnostics). After 20-h incubation period with BrdU, the absorbance was measured at 450 nm/ 600 nm using a microplate reader (ELISA-Reader Zenyth 200st).

### Cell growth and anchorage independent growth

To generate growth curves, PFKFB3-4+5-deficient and src-shRNA HEK-293 cells were seeded (5000 cells/12-well) and every 24 h cells were counted until confluency was reached.

Anchorage independent growth was investigated using a soft-agar test. A total of 5000 cells per 6-well were resuspended in 0.4% agarose in DMEM and were plated on top of a 0.6% bottom agarose DMEM layer. The medium was replenished every 2d. After 14d, colonies were counted in five randomly selected fields per well under x10 magnification.

### Statistics

Data were analyzed with GraphPad Prism software (version 7.0, La Jolla, CA). Group means were compared by a two-tailed Student’s t-test, unless the assumption of normality of the sample distribution was violated. In this case group means were compared by a non-parametric rank sum test. Data are reported as mean ± SEM of at least four independent experiments.

## Results

Previously, we have shown that the PFKFB3 splice pattern is notably different between healthy brain tissue and rapidly proliferating malignant gliomas [20]. We found that PFKFB3-1 (UBI2K5) mRNA concentration was elevated in high grade astrocytomas (not published), whereas PFKFB3-4 (UBI2K4) mRNA expression level was decreased when compared to normal brain tissue [21]. Importantly, the quantitation of PFKFB-4 mRNA involved the recently detected PFKFB3-5 (PFKFB3 splice variant 5) because the C-termini of PFKFB3-4 and PFKFB3-5, which harbor the phosphatase activity, are structurally identical, whereas their N-terminal ends, which accommodate the kinase activity, are different (Fig 1).

To gain more detailed insight about the role of PFKFB3-4 and PFKFB3-5 in glioblastomas, we employed the U87 glioblastoma cell line and investigated the knockdown and overexpression of these splice variants in relation to viability and proliferative capacity of U87 cells as a read out. In parallel, we studied these aspects in non-glial HEK-293 cells, as their PFKFB3 splice patterns for PFKFB3-4 and PFKFB3-5 are similar to that of healthy brain tissue (Fig 7A).

### Knockdown of PFKFB3-4+5 reduces proliferation and cell viability

We used RNA interference (RNAi) to reduce the PFKFB3-4+5 expression in both HEK-293 cells (stable knockdown) and U87 cells (transient knockdown) (Fig 2A). The selective inhibition of PFKFB3-4 and PFKFB3-5 was not possible because the variable exons 1A, 1B and D also occur in several other splice variants. First, we measured the transcript quantity of PFKFB3-4+5 in cells stably and transiently transfected with PFKFB3-4+5 siRNA next to control cells expressing the respective scr-siRNA (Fig 2B). As expected both stable and transient knockdown in HEK-293 and U87 cells with PFKFB3-4+5 siRNA showed a significant reduction of PFKFB3-4+5 transcripts compared to scr-siRNA cells. Consistently, western blot analysis of protein extracts from these cells showed reduced levels of PFKFB3-4 (Fig 2C). Notably, PFKFB3-5 protein seems to be expressed in very low copy number and was not detectable in our hands. To determine whether the decreased expression of PFKFB3-4+5 has an effect on proliferation and/or cell viability of HEK-293 and U87 cells, we performed WST and BrdU assays quantifying the metabolic activity and DNA replication rates of cells, respectively (Fig 2D,E). In both cell lines, knockdown of PFKFB3-4+5 resulted in decreased cell viability and proliferation compared to control cells. Interestingly, the effect appeared more pronounced in HEK-293 cells (Fig 2D).

### Knockdown of PFKFB3-4+5 impinges on cell growth and colony formation

To analyze whether the reduction of PFKFB3-4 and −5 affects the cell number, we quantified stably transfected PFKFB3-4+5 shRNA HEK-293 cells for a period of five days. Similar cell numbers were counted in PFKFB3-4+5-deficient and control samples over a period of the first four days. Interestingly, after five days knock-down of PFKFB3-4+5, a significant reduction of cell number compared to control was observed (Fig 3A,B). Moreover, as glioma cells have the capacity to grow three-dimensionally through neuronal tissues, we sought to interrogate the behaviour of PFKFB3-4+5-deficient HEK-293 cells in soft agar by observing colony formation. Cell colony number dropped by 15 % after 14 days, which may mirror the reduction of the malignant facility of these cells (Fig 3C,D).

**Fig 3.**
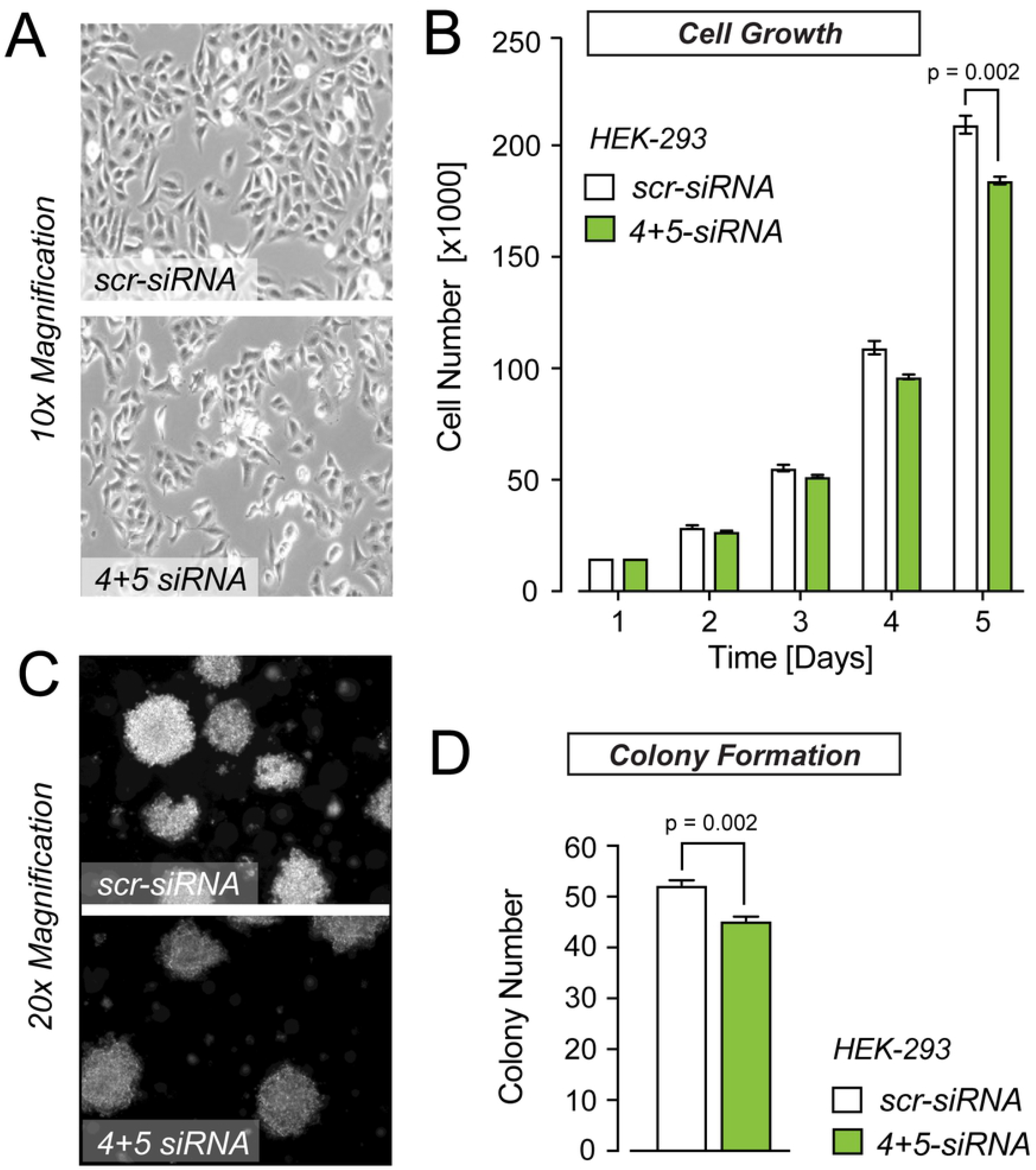
PFKFB3-4+5 knockdown leads to decrease in cell growth and colony formation. (A) Representative brightfield image of scr-siRNA (upper panel) and 4+5 siRNA treated (lower panel) HEK-293 cells cultured in 12-well plates. Image was taken after five days of cell seeding. (B) Quantification of cell growth and colony formation of HEK-293 cells with stably reduced levels of PFKFB3-4+5 compared to cells expressing scr-shRNA. 5000 cells/12-well for each condition. The colony numbers are the mean ± SEM (N=3, n=5). (C) Representative images of soft agar colonies formed by HEK-293 cells (scr-siRNA) and HEK-293 cells with stably reduced PFKFB3-4+5 levels (4/5 siRNA/ The cells (5000 cells) were cultured for 14 days in 6-well plates on soft agar. (D) Quantification of HEK-293 cell colonies from (C) after 14 days in culture. The colony numbers are the mean ± SEM (N=3, n=5).

### Overexpression of PFKFB3-4 and PFKFB3-5 causes opposite effects on cell viability and proliferation

Previously, overexpression of variant PFKFB3-4 C-terminally appended with a biochemical tag (Flag-tag) was shown to reduce both cell viability and anchorage-independent growth of U87 cells [21]. The C-terminal region of PFKFB3-4 encodes the phosphatase moiety of PFKFB3. Hence, it is conceivable that fusion of any tag to this region will disturb the phosphatase function, which in turn may be responsible for these cellular changes. Based on RNAi-mediated effects documented in this study, we hypothesized that overexpression of PFKFB3-4 would lead to an increase in cell viability and proliferation. To test this, we stably overexpressed PFKFB3-4 in HEK-293 and U87 cells. Figure 4A and B show a significant increase of this PFKFB3 variant on transcriptional and translational levels. Indeed, we found that elevated levels of PFKFB3-4 affected proliferation and cell viability positively (Fig 4C,D). In a separate set of experiments, we tested the effects of PFKFB3-5 on these cellular parameters. We followed the same rationale and first validated the transient overexpression of PFKFB3-5 in HEK-293 and U87 cells via qPCR and Western blot analysis (Fig 5A,B). Interestingly, despite the high levels of PFKFB3-5 due to transient overexpression, cell viability and proliferation remained indistinguishable from controls (Fig 5C,D). For this reason, we asked if overexpression of PFKFB3-5 influences the mRNA levels of PFKFB3-1 and PFKFB3-11, splice variants which are constitutively expressed in glioblastoma cells. Transient overexpression of PFKFB3-5 resulted in an increase of PFKFB3-1 in both cell lines, whereas an increase in PFKFB3-11 mRNA was detected exclusively in U87 cells (Fig 5E). Thus, drastic overexpression of PFKFB3-5 impacts the expression level of other PFKFB3 splice variants, indicating their functional interplay.

**Fig 4.**
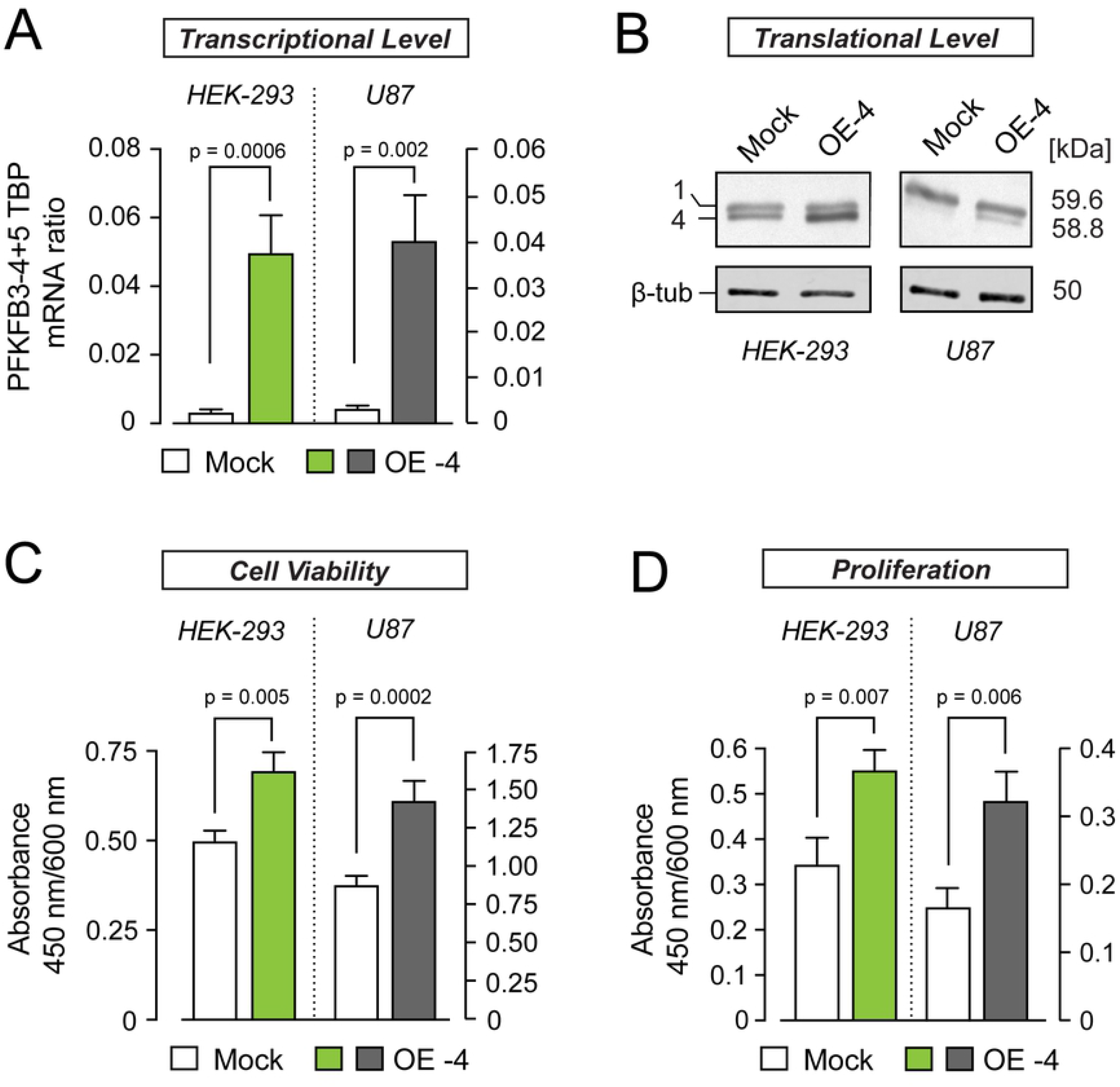
PFKFB3-4 overexpression facilitates cell viability and proliferation. (A) Quantification of PFKFB3-4+5 mRNA levels from HEK-293 (green) and U87 cells (grey) stably and transiently overexpressing PFKFB3-4 (OE −4), respectively. PFKFB3-4 mRNA quantity measured by quantitative PCR was normalized to the amount of TBP mRNA and compared to mock samples. (B) Western blot analysis to confirm the overexpression of PFKFB3-4 with polyclonal PFKFB3 antibody. β-Tubulin served as loading control, 5 μg protein was loaded per lane. (C) Effect of PFKFB3-4 overexpression on cell viability, measured by WST-1 assay. (D) Effect of PFKFB3-4 overexpression on proliferation measured by BrdU-assay. All values represent the mean ± SEM (N=3, n=5).

**Fig 5.**
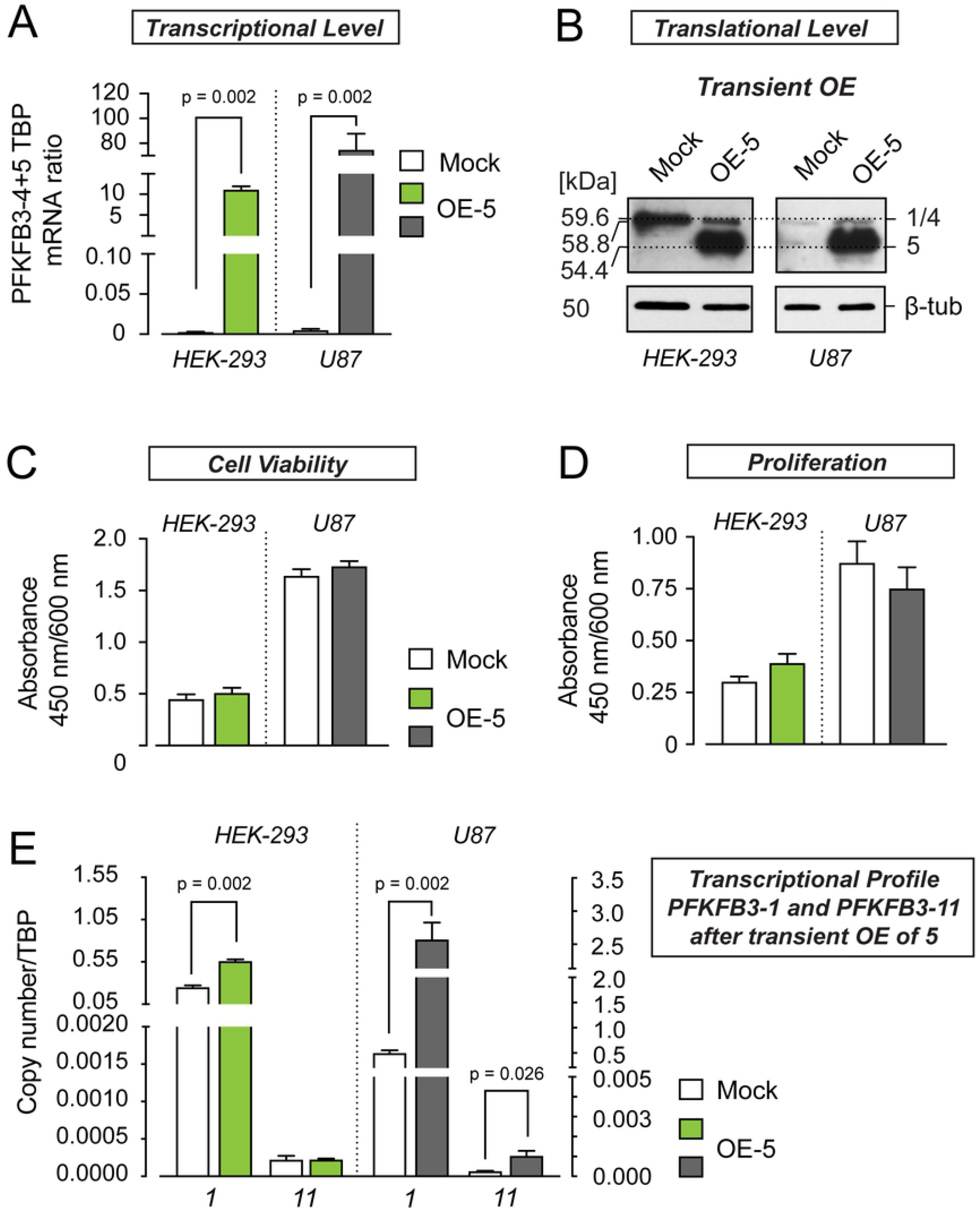
Transient overexpression of PFKFB3-5 has no impact on cell viability and proliferation but changes the transcriptional profile of PFKFB3-1 and 11. (A) Quantification of PFKFB3-5 mRNA levels from HEK-293 (green) and U87 cells (grey) transiently overexpressing PFKFB3-5. PFKFB3-5 mRNA quantity was compared to mock samples and normalized to the amount of TBP mRNA measured by quantitative PCR. (B) Western blot analysis to confirm the overexpression of PFKFB3-5 with polyclonal PFKFB3 antibody. β-Tubulin served as loading control, 5 μg protein were loaded per lane. (C) Effect of PFKFB3-5 overexpression on cell viability, measured by WST-1 assay. (D) Effect of PFKFB3-5 overexpression on proliferation measured by BrdU-assay. (E) Influence of transient overexpression of PFKFB3-5 on the mRNA levels of PFKFB3-1 and PFKFB3-11 compared to mock samples. All values represent the mean ± SEM (N=3, n=5).

To test whether the effects of PFKFB3 splicing on cell viability and proliferation are dosage-dependent, we stably overexpressed PFKFB3-5 in HEK-293 cells (Fig 6A,B). Strikingly, moderate overexpression of PFKFB3-5 has an inhibiting effect on cell viability and proliferation. In contrast, high PFKFB3-5 expression level in transiently transfected HEK-293 cells had no effect on cell viability and proliferation (Fig 5C,D and Fig 6C,D). Noticeably, transcript levels of PFKFB3-1 and PFKFB3-11 appeared unaltered when either PFKFB3-5 or PFKFB3-4 are overexpressed under these conditions (Fig 6E). Taken together, our findings prove specific, dose-dependent effects of PFKFB3 splice variants on the growth capacity of tumor cells.

**Fig 6.**
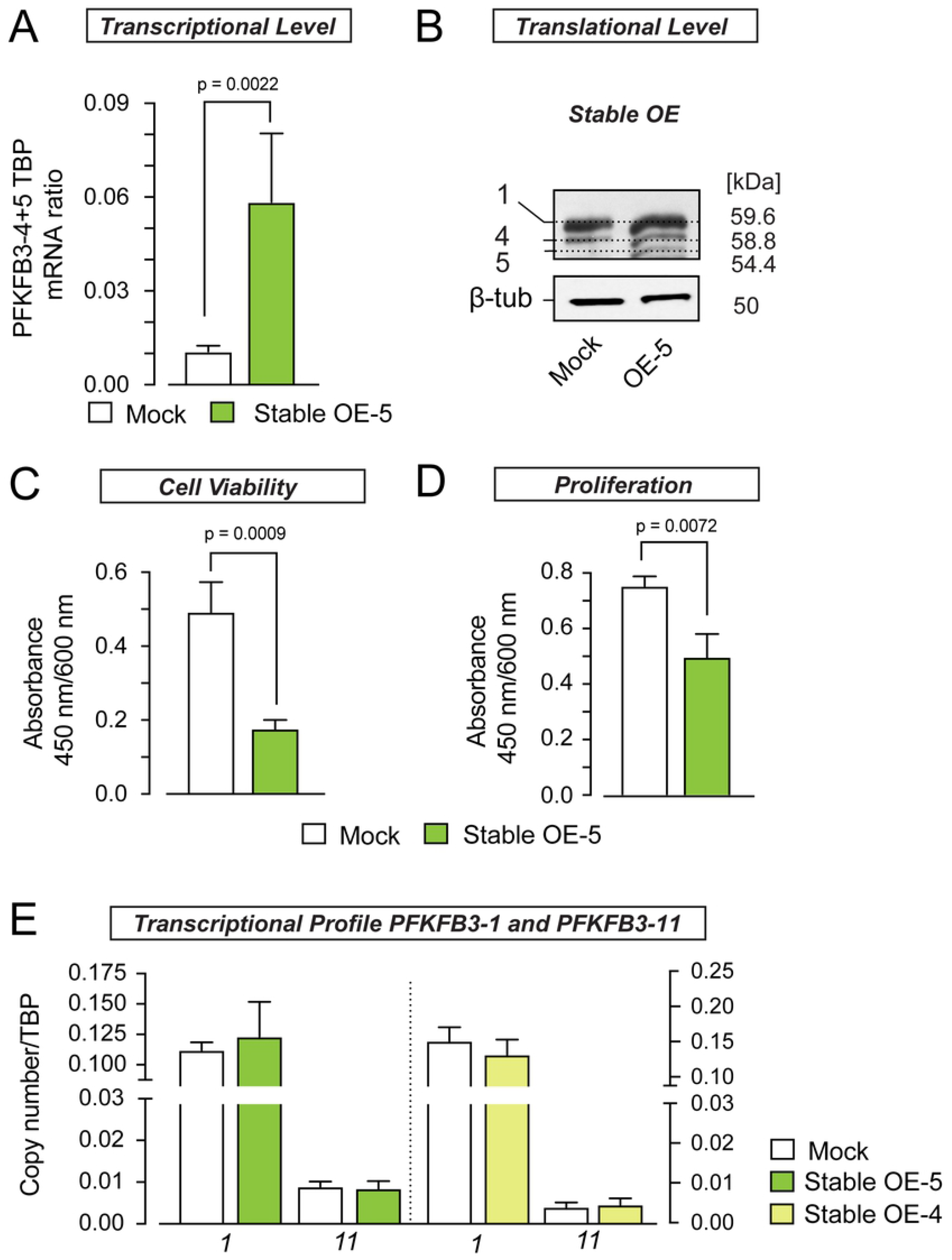
Stable overexpression of PFKFB3-5 leads to decreased cell viability and proliferation while leaving transcriptional profile of PFKFB3-1 and −11 unaltered. (A) Quantification of PFKFB3-5 mRNA levels from HEK-293 cells stably overexpressing PFKFB3-5 (OE −5). PFKFB3-5 mRNA quantity was normalized to the amount of TBP mRNA measured by quantitative PCR and compared to mock samples. (B) Shows western blot analysis to confirm the overexpression of PFKFB3-5 with polyclonal PFKFB3 antibody. β-Tubulin served as loading control, 5 μg protein were loaded per lane. (C) Effect of stable PFKFB3-5 overexpression on cell viability, measured by WST-1 assay. (D) Effect of stable PFKFB3-5 overexpression on proliferation measured by BrdU-assay. (E) Influence of stable overexpression of PFKFB3-5 (light green) and PFKFB3-4 (yellow) on the mRNA levels of PFKFB3-1 and PFKFB3-11 compared to mock samples. All values represent the mean ± SEM (N=3, n=5).

### PFKFB3-5 expression is reduced in glioblastomas (IDH-wildtype)

Contradicting previous reports [21], the data presented here support the idea that PFKFB3-4 exerts no growth-inhibiting effect, while PFKFB3-5 inhibits cell proliferation *in vitro*. This begs the question whether the ratio of PFKFB3-4 to PFKFB3-5 is relevant for neoplastic traits in glioblastomas. To examine the PFKFB3-4 to −5 mRNA ratio, we set up a multiplex PCR to simultaneously measure both transcript species in different cell lines including glioblastoma cells and in glioblastoma patient samples. In glioblastoma cell lines (U87, LN405 and 1321N1), the ratio between PFKFB3-4 to PFKFB3-5 mRNA was significantly shifted toward −4 (U87: 80:1; LN405: 5.4:1 and 1321N1: 5.7:1; Fig 7B,C). Non-glioma cell lines (HEK-293, SH-SYHY) and normal brain tissue samples from the temporal cortex showed a ratio close to 1:1 (HEK-293: 0.65:1; SH-SY5Y: 1.1:1; Fig 7B,C).

**Fig 7.**
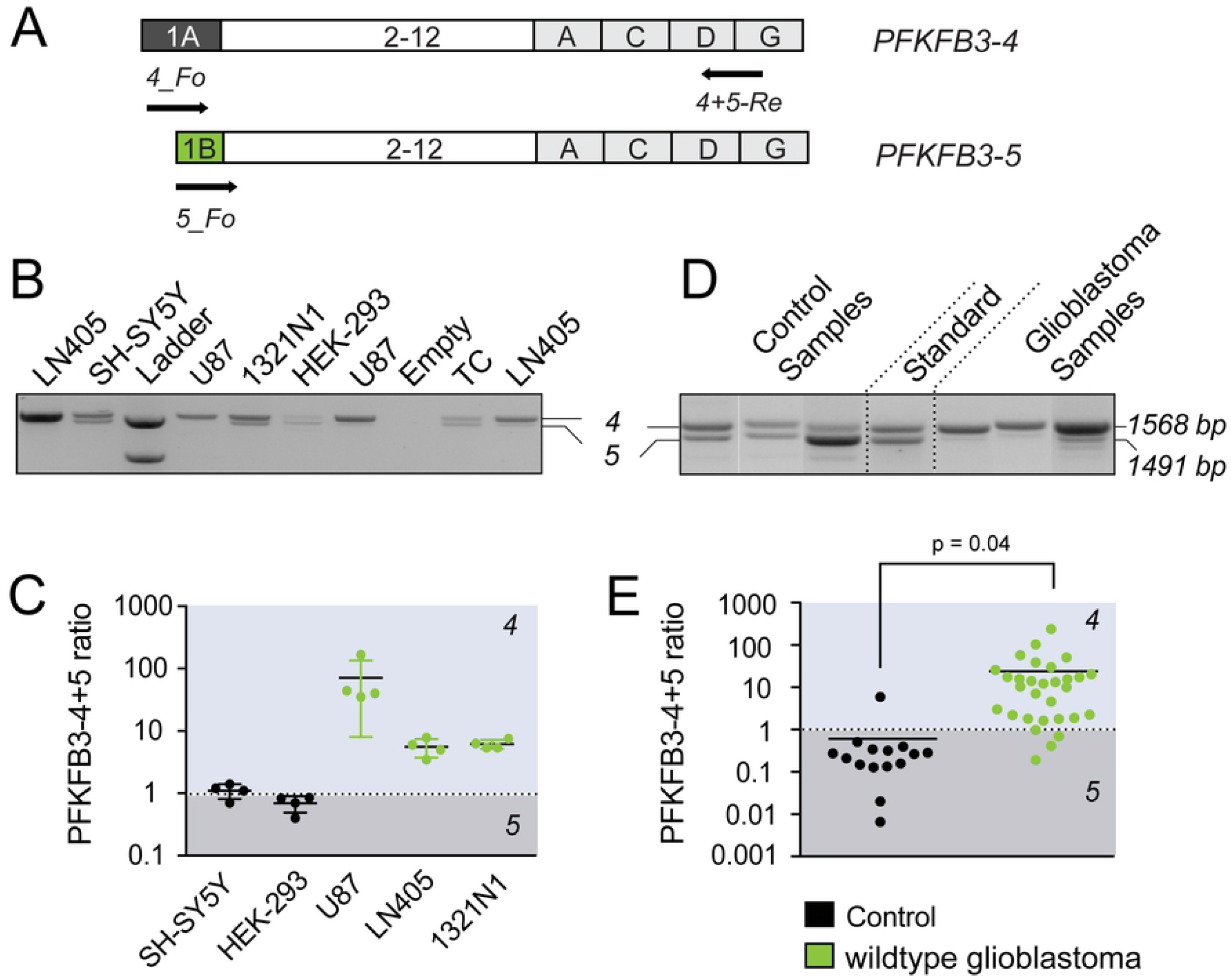
Glioblastoma cell lines and wildtype glioblastomas are signified by high PFKFB3-4 to PFKFB3-5 ratio. (A) Schematic illustration of experimental design of multiplex PCR used to measure the PFKFB3-4 to PFKFB3-5 mRNA ratio in different cell lines (B,C) and brain samples from patients (D,E). (B) Multiplex PCR products from several cell lines and human temporal cortex (TC) were separated by agarose gel electrophoresis. 1500 and 1200 bp fragments were used as standard ladder. (C) Scatter dot blot of PFKFB3-4 to PFKFB-5 mRNA ratios calculated from fluorescence intensities of PFKFB3-4 (1568 bp) and PFKFB3-5 (1491 bp) fragments. (D) Agarose gel electrophorese of multiplex PCR products from three representative normal tissue as control samples and three IDH-wildtype glioblastomas. Equal amounts of mRNA from PFKFB3-4 and PFKFB3-5 (10^7^ copies) were used as a standard. (E) Fluorescence intensities of multiplexed PCR fragments from 15 control and 30 samples from IDH-wildtype glioblastoma patients were used to quantify the ratio of PFKFB3-4 to PFKFB3-5. Similar to cell lines, fast-proliferating glioblastoma samples from patients tend to show higher PFKFB3-4 expression.

Motivated by these findings, we analyzed the PFKFB3-4 to PFKFB3-5 ratio in 30 IDH-wildtype glioblastomas and in 15 normal human brain samples (Fig 7D,E). We found that PFKFB3-4 to PFKFB3-5 ratio in IDH-wildtype glioblastomas (24:1) was about 40-fold higher than in normal brain tissue (1:1.6). Similarly to glioblastoma cell lines, the ratio of PFKFB3-4 to PFKFB3-5 in IDH-wildtype glioblastomas was directed towards splice variant PFKFB3-4. This is in agreement with our findings that PFKFB3-4 promotes proliferation of U87 cells, whereas PFKFB3-5 has an inhibitory effect on cell proliferation. Hence, low PFKFB3-5 expression levels relative to PFKFB3-4 levels seem to confer growth advantage on glioblastomas.

In sum, our data show that PFKFB3-5 may play a decisive role in growth regulation of glioblastomas.

## Discussion

High rates of glycolysis constitute a prerequisite to sustaining the metabolic demands of glioblastomas. The PFKFB3 isozymes have been identified as one of the major metabolic players in glioblastoma however, thus far the functional relevance of PFKFB3 splice variants is only partially understood. The consequences of the different C- and N-terminal structures of PFKFB3 splice variants on their individual functions are unknown. However, the tissue-dependent expression pattern of these splice variants [16] point to their specific functional/regulatory roles in cell metabolism. In humans, at least eleven different PFKFB3 transcripts (PFKFB3-1-11) are known. In glioblastomas only three PFKFB3 transcripts −1, −4 and −11 (former UBI2K4, 5 and 6) were detected, with decreased mRNA levels documented for PFKFB3-4 [20, 21], compared to low-grade astrocytomas and normal brain tissue. Moreover, overexpression of PFKFB3-4 fusion protein blunted cell viability and anchorage-independent growth of U87 cells, and its expression level inversely correlated with the growth rate of several human cancer cell lines [21]. Consistent with the idea that PFKFB3-4 possesses tumor inhibiting features, Fleischer *et al*. identified the loss-of-heterozygosity (LOH) of the *PFKFB3* gene locus, which negatively affects the prognosis of glioblastoma patients [7]. Following this rationale, we expected that knockdown of PFKFB3-4 with siRNA should elevate cell growth. Contrarily, we found that knockdown of PFKFB3-4+5 in U87 and HEK-293 cells results in decreased cell viability and cell proliferation when compared to control samples. Note that the shRNA and siRNA probes used in this study were directed against the C-terminal stretch, the sequence of which is indistinguishable between PFKFB3-4 and PFKFB3-5 (Fig 2A), thus both variants were affected simultaneously. The discrepancy between the growth inhibiting effects induced by PFKFB3-4+5 knockdown, as well as the overexpression of the PFKFB3-4 fusion protein requires a more detailed investigation of PFKFB3-4 in the glioblastoma context, especially with regard to the putative effects of biochemical tag fusion [29].

To mimic (patho)physiological conditions more closely, native PFKFB3-4 was stably overexpressed in HEK-293 and U87 cells. We found increased viability and proliferation of both cell lines compared to control cells with empty vector, indicating growth promoting effects of PFKFB3-4 (Fig 4C,D). This is in line with the blunted growth in PFKFB3-4+5-deficient HEK-293 and U87 cells (Fig 2D,E), strongly arguing against the tumor-suppressive role and rather suggesting tumor-promoting effects of PFKFB3-4. Paradoxically, PFKFB3-4 expression was shown to be reduced in glioblastoma samples versus low-grade astrocytomas and normal brain tissue [21]. This may be reconciled by the fact that the qPCR oligonucleotides target not only PFKFB3-4 mRNA, but also PFKFB3-5 mRNA (Fig 2A), the sequence of which was only recently published in the NCBI database (NM 001323016.2). Therefore, we turned our attention to the investigation of PFKFB3-5 function in glioblastoma. Transient overexpression of PFKFB3-5 in both HEK-293 and U87 cells left viability and proliferation unaltered (Fig 5C,D), but led to a significant increase in PFKFB3-1, an effect not detectable when PFKFB3-4 was overexpressed (Fig 5E, 6E). PFKFB3-1 constitutes the best studied and most abundant PFKFB splice variant in tumor cells known to promote tumorigenic progression [30]. Similar to stable overexpression of PFKFB3-4 we generated a HEK-293 cell line stably overexpressing PFKFB3-5. Strikingly, cell viability and proliferation were decreased in these cells compared to control cells (Fig 6C,D), while PFKFB3-1 and −11 levels remained unaltered (Fig 6E). In conclusion, our data suggest that PFKFB3-5 mediates growth inhibiting effects *in vitro*, while PFKFB3-4 exerts the opposite effect on tumor cell growth. In summary, these results underscore that data derived from exogenous cell systems should be carefully interpreted and the findings should be validated ideally in more native experimental settings. To this end, we quantified the PFKFB3-4 to PFKFB3-5 ratio in different cell lines varying in their proliferation features. Interestingly, astrocytoma cell lines like U87, LN-405 or 1321N1 lines contain more PFKFB3-4 than PFKFB3-5 mRNA, while the PFKFB3-4 to −5 ratio is close to 1:1 in non-glioma cell lines (Fig. 7B,C). Next, we collected glioblastoma and normal brain samples from patients and scored the PFKFB3-4 to −5 ratio. In IDH-wildtype glioblastomas we also found a significant shift towards PFKFB3-4 expression compared to PFKFB3-5 (Fig 7D,E), whereas the PFKFB3-4 to −5 ratio in normal brain tissue was also near 1:1. In conclusion, increased proliferation rates in highly malignant glioblastomas as well as in glioblastoma cell lines might be causally related to the high PFKFB3-4 to −5 expression ratio in which PFKFB3-4 is showing strong growth promoting effects. Our data indicate that, in addition to the well-established pro-proliferating role of PFKFB3-1 [31], also PFKFB3-4 acts as a growth-promoting factor in in glioblastomas.

In order to understand the function of PFKFB3 splice variants their molecular structure has to be contemplated. The enzymatic core of PFKFB3 can be regulated by a variety of different mechanisms [32]. The only structural distinction between PFKFB3-4 and −-5 can be found within the N-terminus, which is typically not post-translationally modified. PFKFB3-5 has a comparably short N-terminus containing only five amino acids, whereas PFKFB3-4 contains 26 amino acids (Fig 1). Based on the crystal structure of PFKFB3 [33] it has been hypothesized that the N-terminus exerts an autoinhibitory effect on PFKFB3 bisphosphatase activity. Bisphosphatase inhibition may thus be relieved to some extent in PFKFB3-5. In accordance with this model a 7-fold higher phosphatase activity was observed for N-terminally truncated versions PFKFB3 [34]. Inversely, it would be interesting to investigate the enzymatic profile of PFKFB3-3, which contains the longest N-terminus amongst PFKFB3 splice variants.

The question remains why glioblastoma cells tend to express less PFKFB3-5. Further, it would be intriguing to study if gliomablastoma cells tend to switch to the expression of splice variants with longer N-termini to ensure an elevation in kinase activity.

Detailed knowledge of putative biochemical differences of PFKFB3 splice variants is scarce. Here, we show that two PFKFB3 splice variants exert different effects on growth rates of cell culture, possibly associated to the structural variation of their N-termini.

Several small molecule inhibitors of PFKFB3 have been developed, although their application to cancer treatment has been limited since tumor cells have developed unique survival strategies to antagonize inhibition of glucose metabolism [35]. More recently, alternative approaches aiming at pharmacological control of PFKFB3’s phosphatase activity were developed and await testing in clinical settings [36, 37]. A *β*-hairpin interaction of PFKFB3’s N-terminus and the phosphatase domain seem to be a structural prerequisite for the autoinhibitory function PFKFB3. Building on this characteristic Macut and colleagues showed that pharmaceutical disruption of this structural element may serve as a handle to increase phosphatase activity [37], which may have the capacity to pave the way towards novel pharmaceutical avenues to treat cancer.

However, the PFKFB3 is embedded in a complex highly regulated metabolic system. In this regard, it should also be mentioned that other PFKFB isoenzymes, especially PFKFB4, shape the adaptation of tumor cell metabolism [38].

In conclusion, we provide experimental and clinical evidence suggesting the significance of a specific PFKFB3 splice variant (PFKFB3-4) as a growth promoting factor in glioblastoma. In addition, here we first report on the role of the novel splice variant PFKFB3-5 in glioblastoma, which contrasts the prevailing growth-promoting function of PFKFB3. Furthermore, our data suggest that the adaptation and survival of tumor cells is shaped by the expression changes of these specific splice variants, a feature that may constitute a first step towards the development of a novel prognostic parameter in glioblastoma.

## Author contributions

MB designed the experiments; RK supported experimental design (patient sample managing and primers); UH and MB performed the experiments and analyzed the data; JPW provided tumor specimens with histological data, KE commented and revised the work; NS and MB prepared figures, wrote and edited the manuscript.

The authors declare no conflict of interest.

## Acknowledgments

This work was supported by the Wilhelm-Sander Stiftung (2004.010.1) and by grant from the Deutsche Forschungsgemeinschaft to NS (FOR2149 P01 [SCHO1791/1-2]). We thank Andrea Boehme for technical assistance as well as Helen Middleton-Price and Tobias Langenhan for discussions.

